# Daptomycin forms a stable complex with phosphatidylglycerol for selective uptake to bacterial membrane

**DOI:** 10.1101/2023.10.01.560395

**Authors:** Pragyansree Machhua, Vignesh Gopalakrishnan Unnithan, Yu Liu, Yiping Jiang, Lingfeng Zhang, Zhihong Guo

## Abstract

Daptomycin is a potent lipopeptide antibiotic used in the treatment of live-threatening Gram-positive infections, but the molecular mechanism of its interaction with bacterial membrane remains unclear. Here we show that this interaction is divided into two stages, of which the first is a fast and reversible binding of the drug to phospholipid membrane in milliseconds and the second is a slow and irreversible insertion into membrane in minutes, only in the presence of the bacteria-specific lipid phosphatidylglycerol, to a saturating point where the ratio of the drug to phosphatidylglycerol is 1:2. Fluorescence-based titration showed that the antibiotic simultaneously binds two molecules of phosphatidylglycerol with a nanomolar binding affinity in the presence of calcium ion. The resulting stable complex is easily formed in a test tube and readily isolated from the membrane of drug-treated bacterial cells, strongly supporting a unique drug uptake mechanism in which daptomycin forms a stable multi-component complex with calcium and phosphatidylglycerol. Revelation of this novel uptake mechanism provides fresh insights into the mode of action of daptomycin and paves the way to new strategies to attenuate resistance to the drug.

## Introduction

Daptomycin (Dap) is a calcium-dependent lipopeptide antibiotic used to treat life-threatening Gram-positive infections.^1, 2^ It exhibits potent bactericidal activity^3, 4^ and was once used as a last-line-of-defence antibiotic against resistant pathogens such as methicillin-resistant *Staphylococcus aureus* and vancomycin-resistant enterococci.^5^ Resistance to Dap is mostly mild with moderate increases in the minimum inhibitory concentration.^6^ However, recent years have witnessed an increasing number of failed treatment due to high-level Dap resistance of clinical pathogens, including *streptococci*,^7, 8^ *Enterococcus faecium*,^9^ and *Corynebacterium striatum*.^10^ Currently, few daptomycin derivatives are available to attenuate the resistance.^11^

The effort to counter the increasing resistance is hampered by our insufficient understanding of the action mechanism of Dap, which has been suggested to involve inhibition peptidoglycan biosynthesis^12, 13^ or membrane depolarization.^14, 15^ Recent studies have provided supporting evidence for the inhibition of peptidoglycan biosynthesis either by disruption of membrane fluidity^16^ or by the formation of a bactericidal complex between the drug and lipid II.^17^ Another unresolved problem is how Dap recognizes Gram-positive bacteria and selectively accumulates in their membrane. Cumulative evidence has shown that this cell specificity is linked to phosphatidylglycerol (PG), a major phospholipid in most bacteria but rare in mammalian cells.^18^ PG was first linked to susceptibility to Dap by the finding that the antibiotic accumulates in membrane microdomains rich in this lipid.^19^ In addition, PG in pulmonary surfactants is able to neutralize Dap, rendering the drug ineffective against community-acquired, bacteria-caused pneumonia.^20, 21^ More importantly, decreased membrane content of PG caused by reduction-of-function mutations in the PG synthase PgsA has been unambiguously shown to be primarily responsible for resistance to Dap, first in the model bacterium *Bacillus subtilis*^22^ and then in clinical pathogens.^23^ When PgsA is inactivated by a single loss-of-function mutation, Dap susceptibility is completely eliminated to result in a very high level of resistance.^24^ Corroborated by *in vitro* liposome model studies,^24^ these findings strongly support a critical role of PG in the recognition of bacterial cells by the antibiotic.

Previous studies have shown that Dap non-specifically binds membrane regardless of the lipid composition^18^ and also interacts specifically with PG in membrane with conformational change.^25–27^ However, the strength of this specific interaction and how it contributes to bacteria-specific uptake of the drug are not clear. In addition, these early investigations reached different conclusion on the stoichiometry of the Dap interaction with PG.^26, 27^ In this study, we investigated the calcium-dependent interaction of Dap with model membranes and found that the mode of interaction is dependent on PG. In absence of PG Dap reversibly binds membrane in a fast process, whereas the drug undergoes a slow, irreversible insertion into membrane when PG is present. Further investigation showed that Dap forms a multi-component complex with calcium and two molecules of PG both *in vitro* and in drug-treated bacterial cells, thus revealing a unique mechanism for the selective uptake of Dap into bacterial membrane.

## Results

### Dependence of Dap uptake on phosphatidylglycerol

Interaction of Dap with membrane is known to increase the fluorescence of its kynurenine residue (Kyn-13, Fig. 1) with a 15 nm blue shift.^28^ We used the kinetic of this fluorescent increase to study the interaction in micelles and found negligible fluorescence increase when the model membrane was comprised of 1, 2-dimyristoyl-*sn*-glycero-3-phosphocholine (DMPC) only (Fig. 2A). When 1, 2-dimyristoyl-*sn*-glycero-3-phosphorylglycerol (DMPG) was added to the phospholipid micelles, the fluorescence increased and reached a plateau in about 10 min. Interestingly, the plateau level is limited by the DMPG content and in most cases is unable to reach a maximum level at which all Dap is absorbed into the model membrane. In addition, the initial rate of the fluorescence change sharply increases with the concentration of both DMPG and Ca^2+^ (Fig. S1). Similar kinetics of Kyn fluorescence was observed in the interaction of Dap with vesicles containing DMPG (Fig. S2), indicating that Dap interacts with PG-containing membrane in the same mode regardless of the model system.

**Figure 1.**
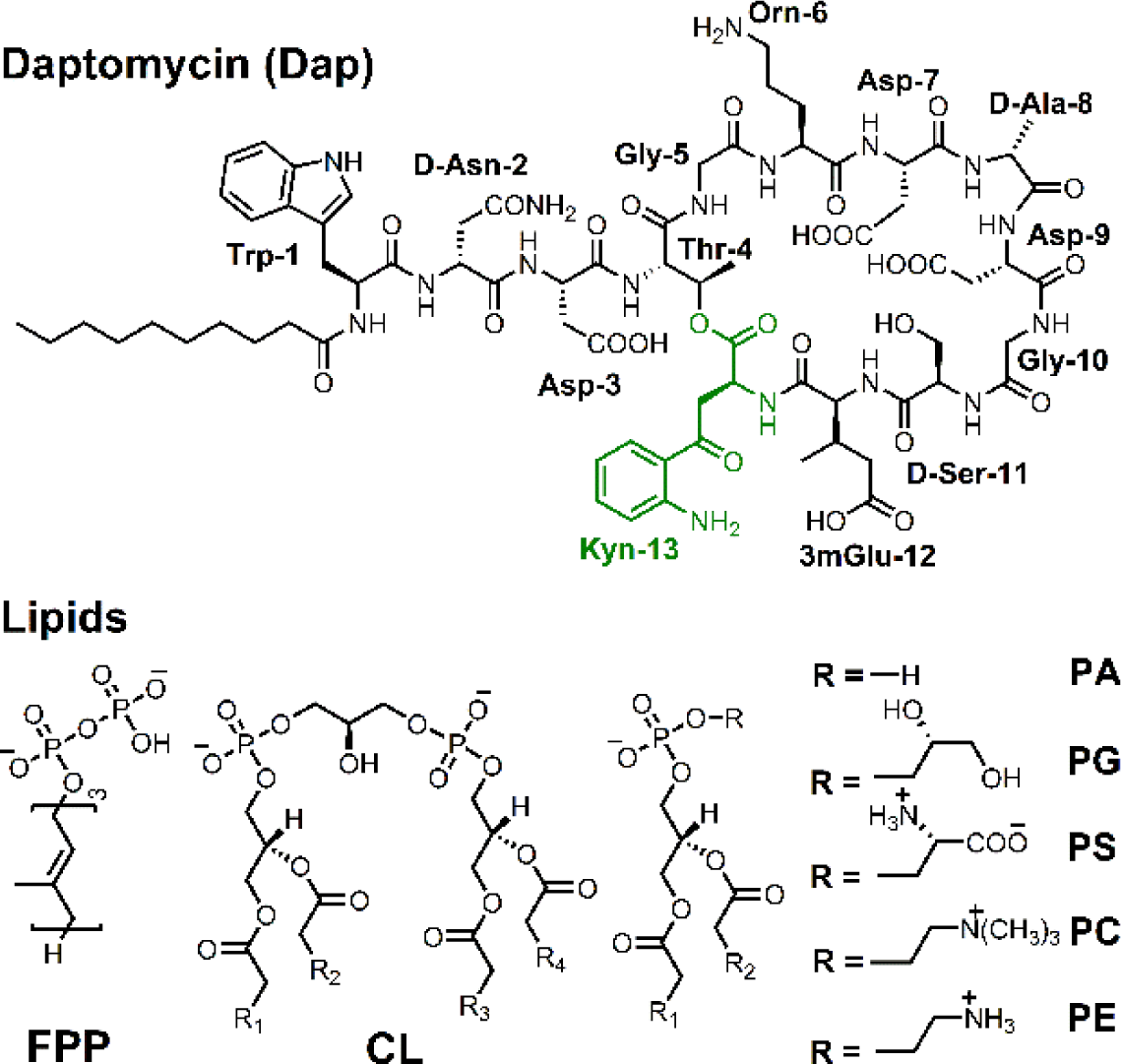
Structure of daptomycin and lipids.

**Figure 2.**
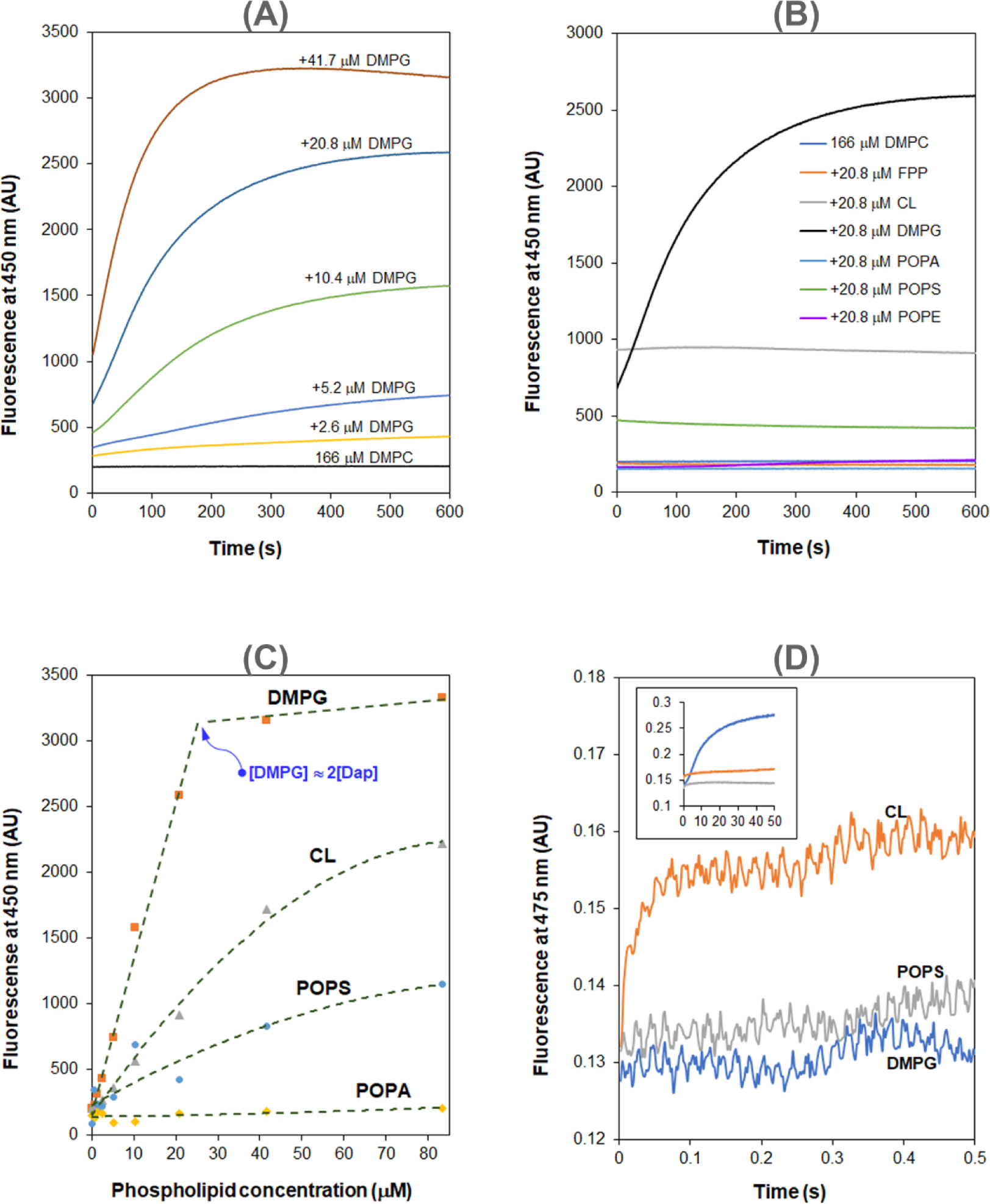
Dap depends on PG in the interaction with membrane. (**A**) DMPG-dependent accumulation of Dap in phospholipid micelles. (**B**) Kinetics of the fluorescence of Dap in interaction with micelles containing different phospholipids. Micelles contained 20.8 μM various phospholipids (11.5%) and 167 μM DMPC. (**C**) Plot of the steady-state Kyn fluorescence *vs*. the content of different negative phospholipids in the micelles. (**D**) A fast Kyn fluorescence increase within 100 ms in the interaction of Dap with CL-containing micelles. The inset shows the Kyn fluorescence change over 50 s; micelles contained DMPG, POPS or CL at 20.8 μM. All the experiments in (**A**)-(**D**) were carried out in 20 mM HEPES (pH 7.57) containing 1.67 mM CaCl_2_; Dap was always added at last at 15 μM. The micelles were prepared from DMPC at 167 μM alone or together with another lipid at a concentration or content indicated in the plots.

Using PG-free micelles, no increase of fluorescence intensity was detected when the model membrane contains the same molar equivalent of other major lipid components in bacteria, including 1-palmitoyl-2-oleoyl-*sn*-glycero-3-phosphoethanolamine (POPE), cardiolipin (CL), 1-palmitoyl-2-oleoyl-*sn*-glycero-3-phospho-L-serine (POPS), 1-palmitoyl-2-oleoyl-*sn*-glycero-3-phosphate (POPA), and FPP (a structural homolog of bactoprenol diphosphate) (Fig. 2B). Nevertheless, the steady-state Dap fluorescence also increases with the content of the lipids, particularly for the negatively charged POPS and CL. Noticeably, this enhanced background fluorescence is accompanied by the same blue shift observed in micelles containing PG (Fig. S3), suggesting that Dap also interacts with the PG-free membrane, but in a mode different from that for the PG-containing membrane. Corroboratively, the steady-state Dap fluorescence is linearly proportional to the DMPG content and reaches a maximum at a concentration about twice that of Dap, but it has a significantly different relationship with the content of POPS, CL, or POPA (Fig. 2C). PG-dependent increase of the steady-state fluorescence was also observed in giant unilamellar vesicles (GUVs).^29^

### Two distinct modes of Dap interaction with membrane

In the interaction of Dap with micelles containing CL, the increase of fluorescence was found to be complete within 100 ms by stopped-flow kinetics (Fig. 2D), demonstrating that this mode of interaction occurs much faster than the slow accumulation of Dap in PG-containing membrane (over minutes, Fig. 2A). This fast process is believed to correspond to the Ca^2+^-dependent recruitment of Dap, via diffusion, to the headgroup region of the phospholipid membrane, where the environment of the Kyn residue becomes more hydrophobic likely due to structural change to cause the Kyn fluorescence enhancement and blue shift. This binding to the membrane surface should be reversible and no Dap is irreversibly inserted into the hydrophobic acyl layer of membrane, since the Kyn fluorescence was not further increased afterwards. Indeed, when Ca^2+^ was sequestrated by EGTA, the bound Dap was completely released back into the bulk solution as indicated by decrease of the Kyn fluorescence to the same level of free Dap control without the lipids (Fig. 3A). In support of this reversibility, the dramatic conformational change identified for Dap bound to the CL micelles was completely reversed to be identical to free Dap after Ca^2+^ sequestration (Fig. 3B).

**Figure 3.**
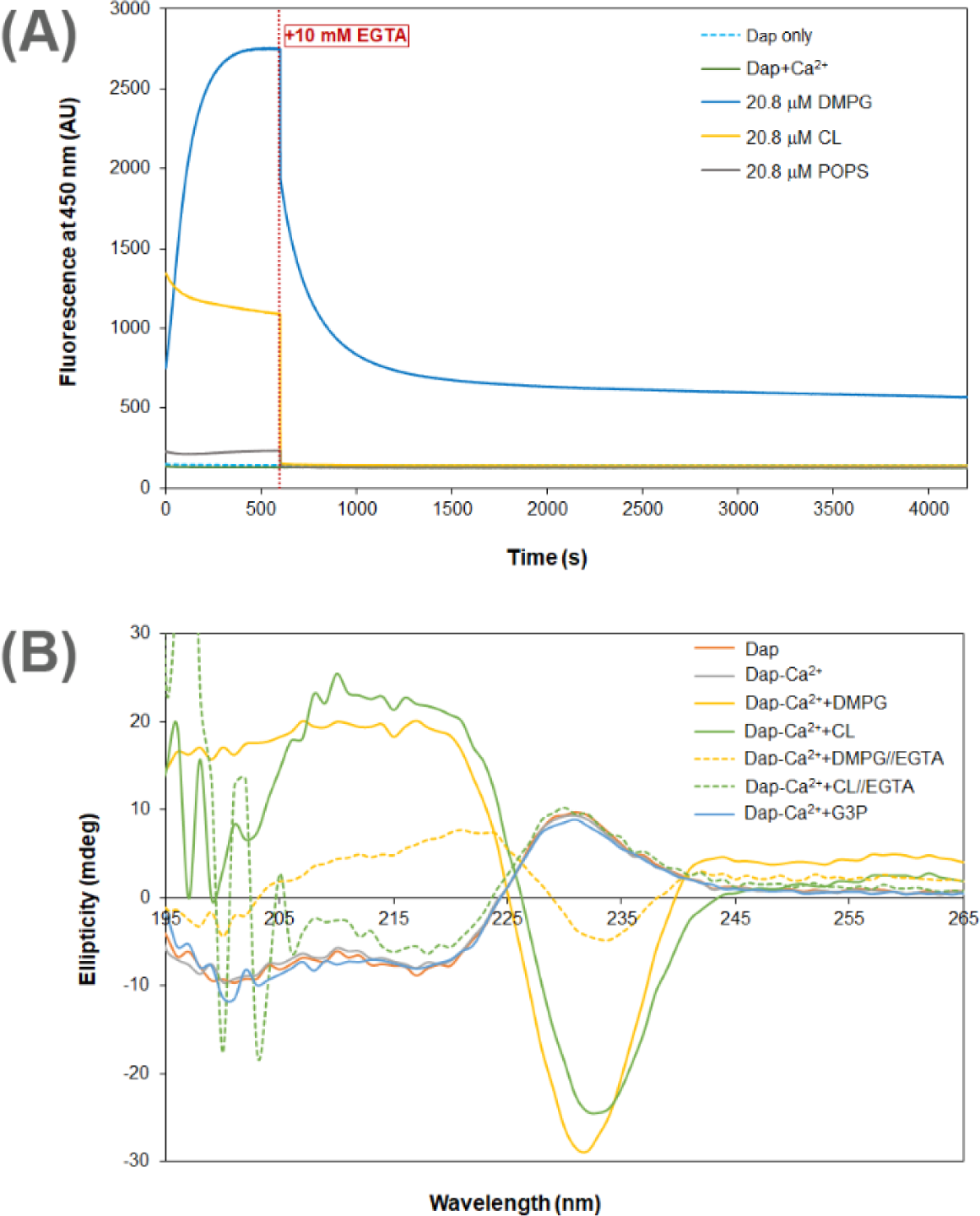
Calcium dependence of the Dap structure and fluorescence in membrane. (**A**) Calcium sequestration decreases the Dap fluorescence in a way dependent on the phospholipid composition. The kinetic fluorescence measurement was performed under the same conditions as in Figure 1(B). The red dotted line denoted the time point when 10 mM EGTA was added. The controls contained Dap at 15 μM Dap in the same buffer with or without calcium. (**B**) Circular dichroism spectra of Dap in different membrane environments. The buffer was 20 mM Tris.HCl, pH 7.57; Dap was 180 μM and Ca(CH_3_COO)_2_ was 1.0 mM; spectra were recorded 30 min after mixing with micelles contained pure DMPG or CL at 720 μM to maximize the amount of bound Dap; in Ca^2+^ sequestration experiments, spectra were recorded 30 min after adding 10.0 mM EGTA. The putative PG headgroup, *sn*-glycerol 3-phosphate (G3P), was 20 mM in attempt to detect its potential interaction with Dap.

By contrast, Dap was inserted irreversibly into the DMPG-containing membrane, because the Kyn fluorescence continuously increased (Fig. 2A) until all Dap was exhausted when the DMPG content was high enough (Fig. 2C). This is supported by the substantial residual Dap fluorescence after EGTA sequestration of Ca^2+^ from the accumulated drug in DMPG-containing micelles (Fig. 3A). This residual fluorescence not only indicates that Dap remains bound to membrane after removal of the metal ion but also demonstrate that Dap is still associated with Ca^2+^ after insertion into membrane. The fluorescence decrease induced by Ca^2+^ sequestration suggests that Dap is buried deeper into membrane when bound by the metal ion. The irreversible insertion of Dap into DMPG-containing micelles is further supported by the conformation of Dap after Ca^2+^ sequestration, which remains significantly different from free Dap albeit with significant change (Fig. 3B). Interestingly, Dap shows a non-identical, similarly shaped circular dichroism spectrum in its reversible and irreversible binding with the CL and DMPG micelles (Fig. 3B), respectively, indicating a small conformational change in the insertion of the surface-bound Dap into the hydrophobic acyl layer of the membrane.

### Ca^2+^-dependent interaction between Dap and DMPG

To assess whether the putative PG headgroup was responsible for the observed irreversible uptake of Dap, G3P was included in the kinetic measurement and found to slightly inhibit the drug uptake at millimolar levels (Fig. S4A). This interaction was not detectable by circular dichroism (Fig. 3B), isothermal titration calorimetry (ITC) or ligand-induced fluorescence change but was found to cause a small fluorescence-based thermal shift in the presence of 20 mM G3P (Fig. S4B). In the NMR titration of G3P into a Dap-Ca^2+^solution at pH 5.40, the α-protons of Trp-1, D-Asn-2, Asp-9 and Kyn-13 were found to shift slightly in the fingerprint region (Fig. S4C). The α-proton signals were assigned by comparing the unique N_α_H-H_α_-H_β_-H_γ_ cross-peak patterns of the amino acid residues from Dap in TOCSY with those obtained for the drug at pH 5.0 in a previous study^30^ (see Fig. S5 for the comparison and Table S1 for the assignments). These results show that Dap indeed binds G3P but the affinity is too low (in high millimolar range) to account for the DMPG-enabled membrane insertion.

Noticeably, Dap was saturated by DMPG at a ratio of 1: 2 in the steady-state fluorescence titration (Fig. 2C), indicating a binding interaction between the two in 1:2 stoichiometry that is consistent with the previous ITC titration.^26^ To determine the binding affinity, Dap at 15 nM was titrated with sub-micromolar DMPG in the presence of Ca^2+^ and its Kyn fluorescence at 454 nm was found to increase sharply at low DMPG concentration and reach a saturating level at high concentration, whereas no fluorescence increase was observed for a control titration with POPS at an elevated Dap concentration of 1.5 μM (Fig. 4). The titration curve was best fitted with a model in which Dap binds two molecules of DMPG with a dissociation constant of K_D_ = 7.2 × 10^-15^ M^2^. DMPG has a critical micelle concentration (CMC) higher than 1.0 μM (Fig. S6) and is homogeneously present in solution in the 0-1.0 μM concentration range under the given conditions. This titration result shows that Dap forms a high-affinity complex with two molecules of DMPG in the presence of calcium ion.

**Figure 4.**
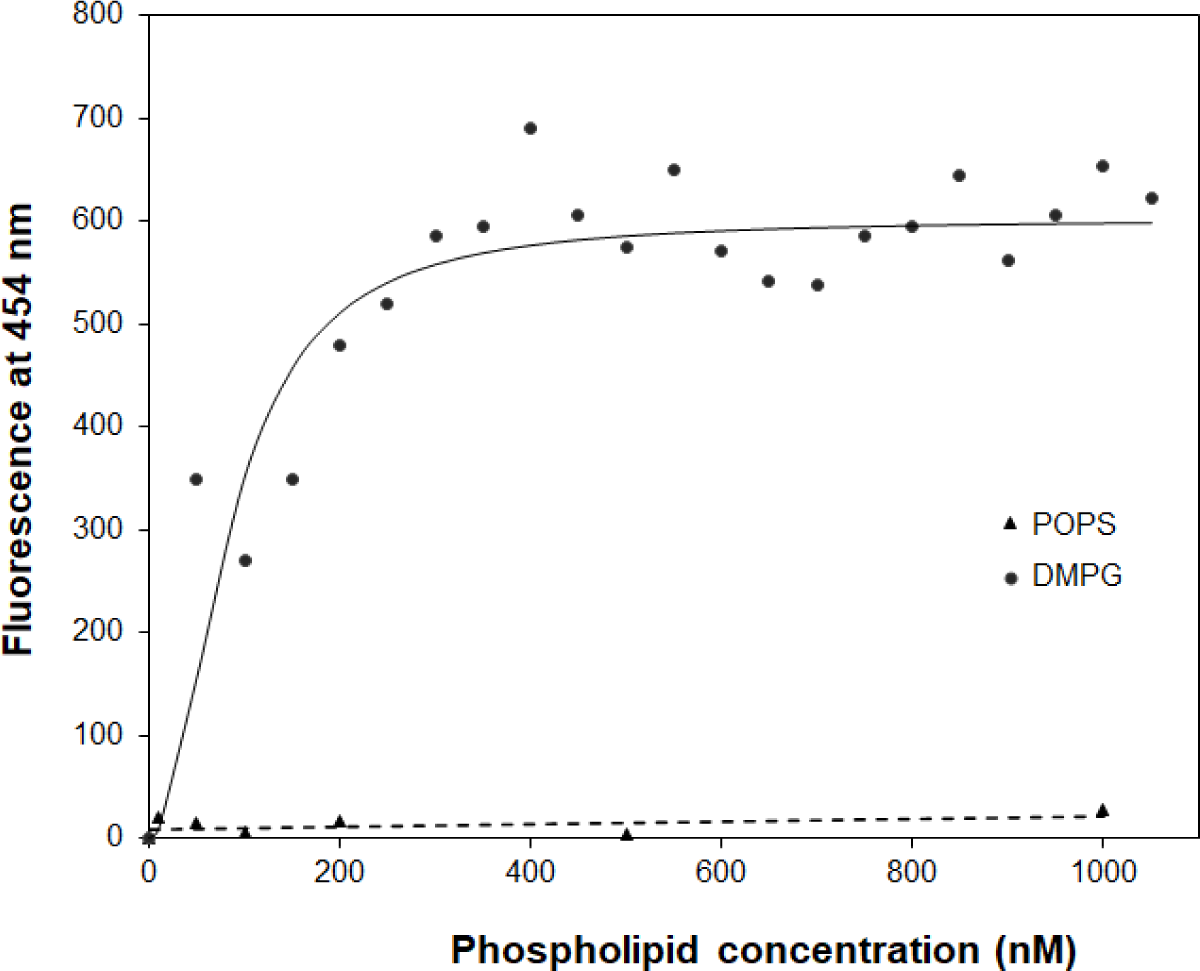
The binding affinity of Dap for DMPG by titration. The titration with DMPG was performed in 20 mM HEPES buffer (pH 7.57) containing 15 nM Dap and 1.67 mM CaCl_2_. The solid line is the fitting curve using the equation F = F_max_ × [DMPG]^2^/(K_D_ + [DMPG]^2^) where F = fluorescence at 454 nm, which is derived from a binding model in which a fluorescent {Dap·2DMPG} complex is dissociated into non-fluorescent Dap and two DMPG molecules under the condition [DMPG] >> [Dap]. The dashed line is the linear trend line for the control titration of 1.5 μM Dap with POPS in the same buffer containing 1.67 mM CaCl_2_.

To obtain the complex, Dap (20 μM or higher) and DMPG were mixed at 1:2 molar ratio in the presence of CaCl_2_ (1 mM or higher) and white precipitate was immediately formed, while mixing DMPG with CaCl_2_ resulted in clear solution (Fig. S7). By comparison, precipitate was formed when POPS or POPA was mixed with CalCl_2_ only, whereas no precipitate was observed when Dap was mixed with other lipids under the same conditions. This comparison suggested that the precipitate was formed from the specific calcium-dependent binding interaction between Dap and DMPG. The same precipitate was also formed by extracting the DMPG-containing micelles after the Dap uptake experiments (Fig. 2A) with chloroform, which was not observed in similar extraction of Dap-bound micelles without DMPG. This precipitate dissolved readily in aqueous solution containing 10 mM EGTA and was moderately soluble in a few organic solvents such as dimethyl sulfoxide (DMSO) and dichloromethane (DCM). In high performance liquid chromatography (HPLC), the dissolved complex was a pure single species in the elution profile (Fig. 5A).

**Figure 5.**
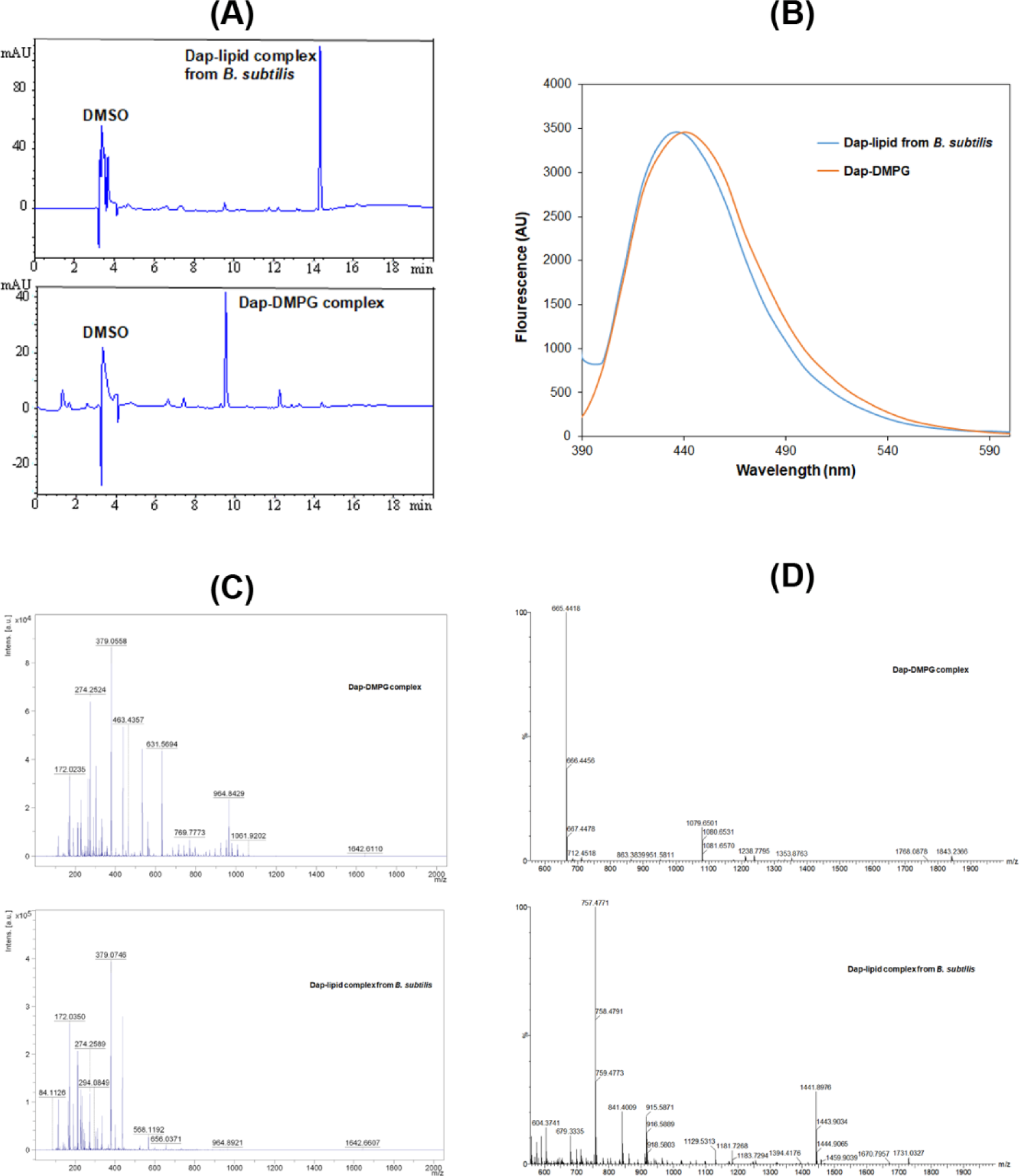
Stable Dap-PG complexes formed *in vitro* and in *Bacillus subtilis*. (**A**) Reverse phase HPLC chromatogram of the complexes. Samples were dissolved in 10% DMSO and 90% water and filtered before injection into a Phenomenex Luna 5 μm C18 column (150 × 4.6 mm). The complexes were eluted by water containing 0.1% trifluoroacetic acid and a linear gradient of acetonitrile from 5-95% over 20 min and detected by ultraviolet absorbance at 254 nm. (**B**) Fluorescence spectra of the two complexes with excitation at 365 nm. The complexes were dissolved in DMSO and the fluorescence of the Dap-DMPG complex was scaled for easy comparison. (**C**) Comparative positive-ion MALDI-ToF mass spectrometry of the two complexes. All the peaks with *m/z* < 500 were from the matrix after comparison with a control. (**D**) Comparative negative-ion high-resolution ESIMS analysis of the two complexes.

### Dap-PG complex isolated from drug-treated bacterial cells

To test whether Dap is complexed with PG in cells, *Bacillus subtilis subsp. Subtilis 168* was grown in Luria broth, treated with Dap at 0. 375 mg/mL and CaCl_2_ at 6.8 mM, and lysed after washing. The cell membrane was isolated from the crude extract by ultracentrifugation and extracted with chloroform. After adding sodium dodecyl sulfate (SDS) to solubilize all precipitated proteins, white precipitate reminiscent of the Dap-DMPG complex was found as suspension in aqueous phase. After dissolved in DMSO, this precipitate was found to contain a pure Dap-containing complex with a retention time different from the Dap-DMPG complex in an HPLC analysis (Fig. 5A). In addition, it was fluorescent with a spectrum characteristic of membrane-bound Dap like the Dap-DMPG complex (Fig. 5B), strongly suggesting that the isolated precipitate was a similar phospholipid complex of Dap. In support of this, Dap was detected with a molecular ion at *m/z* = 1642.66 ([M+Na]^+^, calcd. for C_17_H_101_N_17_O_20_Na: 1642.70) for both the cell-generated Dap-lipid complex and the Dap-DMPG complex in a comparative MALDI-ToF mass spectrometric analysis (Fig. 5C). However, the phospholipid was not detected by MALDI-ToF mass spectrometry. High-resolution electron spray ionization mass spectrometry (HRESIMS) in negative-ion mode turned out to be more sensitive for negative phospholipids, affording DMPG molecular ion at 665.4418 ([M-H]^-^, calcd. for C_34_H_66_O_10_P: 665.4394) for the Dap-DMPG complex (Fig. 5D). For the cell-generated complex, the phospholipid PG (32:0) was deduced from the main peak at *m/z* = 757.4771 ([M+Cl]^-^, calcd. for C_38_H_75_O_10_PCl: 757.4792) in the negative-ion HRESIMS as an adduct with the matrix chloride ion, which could readily be formed in electron-spray ionization.^31^ PG (32:0) likely contains acyl chains derived from two most abundant *iso*-15:0 and *anteiso*-17:0 branched fatty acids in *Bacillus subtilis*,^32^ and its presence in the isolated complex as the major lipid species provides strong support for the specific interaction between Dap and PG. Taken together, all these results consistently indicated that Dap indeed forms a stable complex with PG in bacterial cells, which closely resembles the *in vitro* Dap-DMPG complex.

## Discussion

Daptomycin is well known to target bacterial membrane for its bactericidal effect, but the mechanism of its membrane uptake is poorly understood. By monitoring the kinetics of membrane-induced fluorescence, here we show that the calcium-dependent membrane uptake of Dap is divided into two stages. The first stage is reversible binding to the membrane surface, likely in the headgroup region, which happens very fast and reaches an equilibrium in milliseconds (Fig. 2 and Fig. 3). The Dap binding capacity is dependent on the phospholipid, which is very low for neutral lipids PC and PE but is much higher for negative lipids, particularly cardiolipin (Fig. 2). After this reversible surface binding, Dap is slowly inserted into the membrane in minutes (Fig. 2A and Fig. 3) only in the presence of PG, culminating in the formation of a multi-component complex. This complex contains calcium and two PG molecules with nanomolar affinity (Fig. 4) and is easily formed *in vitro* and readily isolated from drug-treated bacterial cells (Fig. 5). This high-affinity interaction provides an impeccable rationale for the neutralization of Dap by the PG-rich pulmonary surfactants.^20, 21^ Moreover, the complete dependence of the membrane insertion on PG also explains why Dap selectively attacks Gram-positive bacteria without affecting mammalian cells, because PG is a major phospholipid in bacterial membrane but is a minor component in mammalian membrane.

Using circular dichroism spectroscopy, Dap was structurally examined in every step of its interaction with membrane. In bulk aqueous solution, it takes a conformation with a characteristic positive peak at 230 nm and undergoes a large conformational change when bound to the headgroup region of membrane without PG, as evidenced from the dramatic change in circular dichroism (Fig. 3B) that is consistent with previous investigations.^25, 27^ It may not bind Ca^2+^ in the bulk solution but requires the ion to bind the phospholipid headgroups, due to the observed release of the bound drug to the bulk solution with concomitant conformational reversal in the Ca^2+^ sequestration experiments (Fig. 3). In this surface-bound state, Dap is unable to insert into the acyl layer in membrane, suggesting that its decyl tail is folded back into the hydrophobic part of the cyclodepsipeptide ring. In addition, due to the specificity for negative phospholipids (Fig. 2B and 2C), this reversible binding of Dap likely involves both a non-specific Ca^2+^-mediated ionic interaction and a specific interaction with the remaining part of the headgroups.

In the irreversible insertion into membrane, Dap undergoes a subtle conformational change from its surface-bound state to the inserted state, as seen from the small difference in their circular dichroism spectra (Fig. 3B). When Ca^2+^ is sequestrated, Dap undergoes another major conformational change from its inserted state to a new structure with circular dichroism features distinct from all other structures, supporting that the drug is still bound to Ca^2+^ in its membrane-inserted state and that its decyl tail is fully extended and firmly embedded in the hydrophobic acyl layer of membrane. Besides binding Ca^2+^, this inserted Dap should form a complex with PG in 1:2 ratio as indicated by titration of Dap with DMPG in both micromolar (Fig. 2C) and nanomolar range (Fig. 4), which was formed easily *in vitro* and readily isolated from drug-treated *Bacillus subtilis* (Fig. 5). This complex is structurally similar to the tripartite Dap-PG-undecaprenyl lipid complex proposed to be responsible for the bactericidal effect of the drug.^17^ At present, it is not clear whether the Dap-Ca^2+^-2PG complex contains one or more calcium ions.^33, 34^ For simplicity of discussion, only one calcium ion is assumed to involve in the binding interaction to result in a quaternary Dap-Ca^2+^-2PG complex.

Taking all the structural changes into consideration, we propose a mechanism for the Ca^2+^-dependent uptake of Dap into bacterial membrane as shown in Fig. 6. In the reversible binding, Dap with a relatively random conformation is quickly bound to the headgroups of phospholipids likely through Ca^2+^-dependent electrostatic interaction, accompanied by a large conformational change. With a hidden decyl tail, the surface-bound Dap is unable to insert into the membrane, even for the drug molecule bound by the PG headgroup. To enable membrane insertion, we propose that the bound Dap has to form a quaternary complex resembling the post-insertion Dap-Ca^2+^-2PG complex, which is different from the latter in having a hidden rather than a fully extended decyl tail and is less stable. Subsequently, this pre-insertion complex undergoes structural change to expose the decyl tail of Dap and allows it to interact with the acyl chains of the two PG molecules to finish the insertion process, driven by the free energy to form the more stable post-insertion complex.

**Figure 6.**
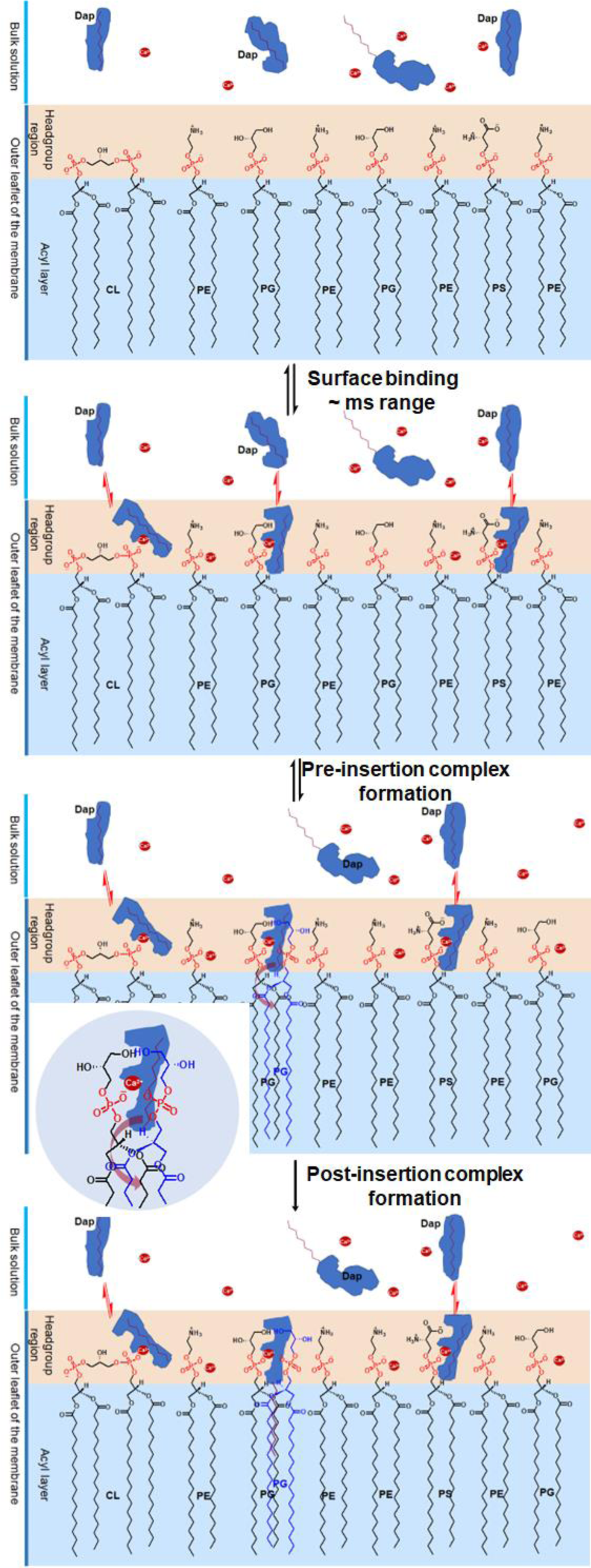
Proposed mechanism for the two-phased uptake of Dap into bacterial membrane. In the first phase, Dap reversibly binds to negative phospholipids with a hidden tail in the headgroup region, where it combines with two PG molecules to form a pre-insertion complex. In the second phase, the hidden tail unfolds and irreversibly inserts into the membrane. The inset shows the headgroup of the pre-insertion complex with the broad arrow showing the direction for the unfolding of the hidden tail. The red dots denote Ca^2+^.

From this uptake mechanism, we can now understand how Dap recognizes bacteria and selectively accumulates in their membrane in a clinic setting. After absorption into the circulation, Dap is preferentially bound to the headgroup region of bacterial membrane due to the high content of PG and CL,^35^ very quickly in milliseconds, although it is also bound to mammalian cell surface in a smaller amount per cell due to the very low content or absence of PG or CL in plasma membrane,^36^ but in substantial total quantity due to the overwhelmingly larger amount of host cells. However, Dap absorbed on bacterial surface is continuously inserted into the acyl layer via formation of complex with PG in a time scale of minutes, whereas little irreversible insertion of Dap occurs on mammalian membrane due to the low content of PG while the bound Dap is continuously released to the circulation as the drug is depleted by the bacteria. The proposed requirement of the pre-insertion quaternary complex increases the threshold of PG content for the membrane insertion to happen and thus makes it less likely on the surface of mammalian cells that contain PG at a low level in the membrane. Consequently, most circulating drug is selectively inserted and accumulated in bacterial membrane within a short period of time where it exerts bactericidal effect via a mechanism still in debate.^12–17^

In the Dap uptake, PG plays the dual role of recruiting the drug to bacterial cell surface and enabling the irreversible insertion of the drug, of which both processes are critically affected in velocity and quantity by its membrane content. This PG dependence not only provides the basis for the reliance of Dap susceptibility on PG,^22^ but also is fully consistent with the finding that a decrease in membrane PG content inevitably leads to resistance to Dap, no matter the decrease is due to the gain-of-function mutations of MprF in the domain responsible for lysinylation of PG,^37–40^ or loss-of-function mutations of the PG synthase PgsA,^22, 41^ or gain-of-function mutations of the cardiolipin synthase (Cls) to increase conversion of PG to CL.^42–45^ Understandably, when the PG biosynthesis is abolished due to complete inactivation of the PgsA activity, the Dap susceptibility is completely lost to result in a very high level of resistance.^23, 24^ However, a decreased PG content is found in only a fraction of resistant Enterococcus strains containing mutations in the LiaFSR operon and the PG metabolic enzymes GpdD and Cls in clinical isolates of Enterococci,^46, 47^ suggesting that there exist PG-independent mechanisms to cause Dap resistance. In this connection, it is noteworthy that the PG content also appears not to be involved in the resistance caused by up-regulated expression of *dltABCD* operon that increases the production of teichoic acids and their D-alanylation.^48, 49^

In summary, we have shown that Dap is first reversibly bound to the membrane surface and then irreversibly inserted into the acyl layer by forming a stable complex with PG. Interestingly, the stable complex may exchange one PG molecule with an undecaprenyl lipid to form the previously proposed bactericidal Dap-PG-undecaprenyl lipid tripartite complex to play a critical role in the antibacterial activity of the antibiotic. In addition, this stable complex also forms the basis of a unique mechanism for the bacterial uptake of Dap, in which the negative phospholipid PG plays a pivotal role to affect both the susceptibility and resistance to Dap. Nevertheless, it remains unknown how the structural components of the drug affect nonspecific binding to the surface headgroup layer and specific binding with PG to influence the speed and dose of drug uptake. Elucidation of these structure-activity relationships may enable rational design of Dap analogues to counter the increasing drug resistance.

## Materials and Methods

### Materials

Daptomycin (Dap) was purchased from MedChem Express. 1, 2-Dimyristoyl-*sn*-glycero-3-phosphocholine (DMPC), cardiolipin sodium salt (CL) from the bovine heart, farnesyl pyrophosphate ammonium salt (FPP), calcium chloride and calcium acetate were purchased from Sigma. The putative headgroups *sn*-glycerol-3-phosphate bis (cyclohexyl ammonium) salt (G3P) was also purchased from Sigma. Tris base was purchased from Fisher Bioreagents. 1, 2-Dimyristoyl-*sn*-glycero-3-phosphate sodium salt (DMPG) was purchased from Abcam. 1-Palmitoyl-2-oleoyl-*sn*-glycero-3-[phospho-L-serine] sodium salt (POPS) and 1-palmitoyl-2-oleoyl-*sn*-glycero-3-phosphate monosodium salt) (POPA) were purchased from Avanti Polar Lipids. All lipids or chemicals were used directly without further treatment.

### Kinetics of the Kyn fluorescence change in micelles

Lipid stocks were prepared in 20 mM HEPES buffer with the pH adjusted to 7.57 at a concentration of 1.0 mM for DMPC and 0. 50 mM for other lipids including DMPG, POPA, POPE, POPS, CL, and FPP. In a typical kinetic measurement, a lipid micelle solution was prepared by mixing DMPC with or without another lipid in 20 mM HEPES (pH 7.57) and was sonicated for at least 4.5 h (Ultrasonic Cleaner, 37 kHz, 100%). After the micelle solution was added calcium chloride and Dap at the last, the kinetic of the Kyn fluorescence at 450 nm was immediately monitored in an Edinburgh FS-5 spectrofluorometer for 600 s with a time interval of 0.1 s and an excitation wavelength of 365 nm. The total volume of the solution was 300 μL (of which only 200 μL was added to the cuvette for reading) containing 167 μM DMPC with or without another lipid at a varied concentration, 1.67 mM CaCl_2_, and 15.0 μM Dap in 20 mM HEPES (pH 7.57). After the kinetic measurement, an emission spectrum was recorded from 380 nm to 700 nm.

To study the effects of putative lipid headgroups on the fluorescence kinetics, their stocks were prepared at 50 mM with the pH adjusted to 7.57. The putative headgroup was then added at a pre-set concentration to the micelle solution for the kinetic measurements. In these measurements, final DMPC concentration was set at 80 μM and final DMPG concentration was set at 10 μM (or 11.5%), while all other components and the instrumental parameters remained unchanged.

In the calcium sequestration experiments, an EGTA stock was prepared at 42.85 mM and dissolved in HEPES by NaOH and adjusted to pH 7.57 by HCl. It was then added at a saturating concentration (10 mM) to the micelle solution at the end of the 600 s kinetic measurement and the fluorescence kinetics was monitored at 450 nm for another 10 min or a longer time (up to 70 min). The micelle solution contained 167 μM DMPC with or without another phospholipid, namely DMPG, or CL, or POPS at 20.8 μM (or 11.5%). All other conditions were the same as in other kinetic measurements. Similar experiments were also performed with a different combination of phospholipids such as 167 μM DMPC and 10.4 μM DMPG.

### Kinetics of the Kyn fluorescence change in vesicles

Phospholipid vesicles were prepared in a varied ratio from a DMPC stock and the stock of another phospholipid, namely DMPG, CL or POPS, of which both were in chloroform at a concentration of 500 μM and 125 μM, respectively. After mixing, the solvent chloroform was evaporated by a Rotavap under reduced pressure at 37°C for 15 min, to form concentric rings at the bottom of the round bottom flask. The phospholipids were further dried under vacuum for 2 h and then hydrated overnight in 20 mM HEPES (pH 7.57). The hydrated phospholipid film was subjected to five freeze-thaw cycles, in each of which the vesicles were frozen in liquid nitrogen for 1 min, followed by 5 min thawing and 10 s vortex. Finally, the vesicle suspension in a 1 ml syringe was pushed through a 100 nm filter in an Avanti polar extruder with filter supports. Vesicles went through at least 10 passes and the extruder was warmed on a hot plate to ease the passing of vesicles. The vesicle suspension was diluted to the preset concentration of the phospholipid and used to interact with Dap for kinetic monitoring of the Kyn fluorescence similar to the measurements using the micelles.

### Dap conformational change by circular dichroism spectroscopy

Circular dichroism (CD) was used to characterize the structure of Dap in its interaction with calcium, G3P and lipid micelles under various conditions using a Chirascan™ Circular Dichroism Spectrometer (Applied Photophysics). The samples were prepared to contain 0.30 mg/ml Dap from stocks adjusted to pH 7.57, to avoid pH change after mixing of different components. To reduce background, 20 mM Tris buffer (pH 7.57) was used instead of the HEPES buffer and calcium acetate was used at 1.0 mM to replace calcium chloride used in the fluorescence experiments. Each sample was scanned 10 times in a 1 mm Hellma Quartz cuvette from 180 to 270 nm with 1 nm bandwidth, 1 nm step size and 0.5 s per point (approximate 73 seconds). The CD spectrum was then averaged from the 10 scans using the Chirascan software after auto-subtraction with the background signal of a blank buffer.

For Dap interaction with model membranes, the micelle solution was prepared from lipid stocks adjusted to pH 7.57 to contain one or two lipids at preset concentrations, sonicated for 4 h, and then added 1.0 mM calcium acetate and 0.30 mg/mL Dap for incubation at room temperature for at least 15 min before the CD spectra were taken. To maximize the drug reversibly bound to the micelle surface, pure CL micelle solution was used at 720 μM (higher concentration led to precipitation) to bind Dap in the presence of 1 mM calcium acetate for CD spectroscopy. Similarly, pure DMPG micelles at 720 μM was used in the presence of 1 mM calcium acetate to ensure complete membrane insertion of Dap for structural characterization by CD spectroscopy. To characterize the conformation of membrane-bound Dap after calcium removal, EGTA in 20 mM HEPES at pH 7.57 was added at a saturating concentration of 10 mM to the solution of Dap bound to micelles of either pure CL or pure DMPG at 720 μM, incubated at room temperature for at least 30 min and then subjected to CD spectroscopy.

### Interaction of Dap with G3P by NMR spectroscopy

For the NMR analysis, stocks of calcium chloride, G3P and Dap were prepared in 20 mM HEPES with 90% D_2_O (v/v) and pH/pD adjusted to 5.40. They were used to prepare the samples (in 1000 μL,) that contained 1.0 mM Dap, 1 mM CaCl_2_ and G3P at a varied concentration at 0 mM, 1.0 mM, 4.0 mM, 8.0 mM, 12.0 mM, 16.0 mM and 20.0 mM in the same deuterated buffer. Controls containing 1.0 mM Dap or 20 mM G3P were also prepared for the NMR experiments in the same buffer. The 1D ^1^H-NMR spectrum and standard 2D spectra including TOCSY (spin lock system was 75 mS), COSY and HMBC were recorded for each sample on an 800 MHz Varian spectrometer at room temperature (23°C). All spectra were analyzed using Mestrenova and the cross-peaks in the fingerprint region were assigned by comparing the N_α_H-H_α_-H_β_-H_γ_ correlation patterns of the amino acid residues of Dap in TOCSY with those obtained for the drug at pH 5.0 in a previous study.^30^ The assignments are shown in Table S1 and the TOCSY comparison is shown in Fig. S5.

### Critical micelle concentration (CMC) of DMPG and titration of Dap with DMPG in the nanomolar range

The CMC of DMPG was determined using a reported method.^50^ Briefly, pyrene (∼ 20 mg) was crushed to powder and added to 5.0 mL to obtain a 1.0 mM stock of DMPG in 20 mM HEPES buffer (pH 7.57). The mixture was sonicated for 4.5 h, filtered to remove undissolved pyrene, and then diluted in the appropriate medium (either pure water or 20 mM HEPES, pH 7.57) to contain DMPG at a varied concentration from 10 nM to 10 μM. All samples were added calcium chloride to a final concentration of 1.67 mM and their emission spectra from 340 nm to 600 nm were recorded on an FS-5 Edinburgh spectrofluorometer with an excitation wavelength at 330 nm and a bandwidth of 3 nm. The ratio of the fluorescence intensity at 473 nm and 484 nm was plotted against the DMPG concentration to determine CMC, as shown in Fig. S5.

For the fluorescence titration at low concentrations, each sample was prepared in 300 μL to contain 1.67 mM CaCl_2_, 15 nM Dap and DMPG at a varied concentration ranging from 15 nM to 10 μM in an appropriate medium (pure water or 20 mM HEPES, pH 7.57). DMPG was added from a stock of DMPG at 5.0 μM in 20 mM HEPES (pH 7.57) while Dap was added from a stock at 150 nM. These samples were then recorded for their emission spectra in the range of 380 nm to 700 nm on an FS-5 Edinburgh spectrofluorometer with the excitation wavelength set at 365 nm. The bandwidth was set at 3 nm for both the excitation and emission. The emission value at 450 nm was then plotted against the DMPG concentration (as shown in Figure 4A). A control titration was performed similarly using 1.5 μM Dap against POPS in the concentration range of 0 – 1.0 μM.

### Isolation of Dap-lipid complex from drug-treated *Bacillus subtilis*

*Bacillus subtilis* 168 was grown in 500 mL LB for 12 h at 37°C inoculated with 1 mL overnight LB culture from a single colony, harvested, resuspended in 10 mL fresh LB, and then added 0.375 mg/mL Dap and 0.75 mg/mL CaCl_2_ for incubation at 37°C for another 2 h. The drug-treated cells were pelleted, washed once with 10 mL 0.9% NaCl solution, resuspended in 10 mL 1 mM NaCl and lysed by sonication (1s sonication, 2 second pause, Instrument: JY92-II) for 1 h over ice. The lysate was centrifuged to remove the cell debris and the supernatant was ultracentrifuged at 125,000 × g at 4°C for 1 h in Hitachi CP80WX Preparative Ultracentrifuge. After removing the supernatant, the pellet was suspended in 2 mL of 0.9% NaCl solution and extracted with 2 mL chloroform to observe a layer of white precipitate at the phase interface. At this point, 1 mL of 0.1 M SDS solution was added to the aqueous phase and the mixture was vortexed for 1 min. 3 mL of DMSO was then added to quench the bubbles and the aqueous phase appeared cloudy. The aqueous layer was separated and centrifuged at high speed to obtain a small amount of white pellet, which exhibits fluorescent emission typical of membrane-absorbed Dap at 365 nm excitation after suspension in water or being dissolved in DMSO. In a parallel control experiment starting from the same amount of cell culture treated with CaCl_2_ but without Dap, the aqueous phase was clear without any precipitate after addition of SDS, indicating that SDS dissolved all protein precipitates formed from the chloroform extraction. Like the Dap-DMPG complex, the final precipitate or pellet obtained from the drug-treated cells was soluble in DMSO and partly soluble in CH_2_Cl_2_ and was subject to analysis by reverse-phase HPLC, MALDI-ToF mass spectrometry and high-resolution ESIMS.

## Abbreviations

CL: cardiolipin
Dap: daptomycin
Kyn: kynurine
FPP: farnesyl pyrophosphate
PC: phosphatidylcholine
DMPC: 1,2-dimyristoyl-*sn*-glycero-3-phosphocholine
PG: phosphatidylglycerol
DMPG: 1,2-dimyristoyl-*sn*-glycero-3-phosphorylglycerol
PA: phosphatidic acid
POPA: 1-palmitoyl-2-oleoyl-*sn*-glycero-3-phosphate
PE: phosphatidylethanolamine
POPE: 1-palmitoyl-2-oleoyl-*sn*-glycero-3-phosphoethanolamine
PS: phosphatidylserine
POPS: 1-palmitoyl-2-oleoyl-*sn*-glycero-3-phospho-L-serine
G3P: *sn*-glycerol 3-phosphate
oPS: *O*-phosphoserine
oPC: *O*-phosphocholine
oPE: phosphorylethanolamine or *O*-phosphoethanolamine

## Funding information

This work was supported by GRF16102121 from the Research Grants Council of Hong Kong SAR and the grant 21877094 from the National Natural Science Foundation of China.

## Author contributions

Conceptualization: Z.G. Methodology: P. M., V. G. U., Y. L., Y. J., L. Z. Investigation: P. M., V. G. U., Y. L, L. Z. Supervision: Z. G. Writing—original draft: Z. G., P. M. Writing—review & editing: Z. G., P. M., V. G. U., Y. L., Y. J.

## Competing interests

Authors declare that they have no competing interests.

## Notes

### Competing Interest Statement

The authors have declared no competing interest.

### Summary of Updates

Figure 2, 3 and 6 revised. Supplemental file updated.

