## Supplementary information for "Daptomycin forms a stable complex with phosphatidylglycerol for selective uptake to bacterial membrane"

**Contents:**

Fig. S1. Dependence of the Dap uptake on both DMPG and calcium. S2
Fig. S2. Enhancing effect of phosphatidyl glycerol (PG) on the Dap uptake to vesicles. S3
Fig. S3. Fluorescence enhancement and blue-shift caused by negative cardiolipin (CL) and phosphatidyserine (PS). S4
Fig. S4. Interaction of Dap-$Ca^{2+}$ with glycerol-3-phosphate (G3P). S5
Fig. S5. Highly similar $N_{\alpha}H$-$H_{\alpha}$-$H_{\beta}$-$H_{\gamma}$ spin systems in Dap at pH 5.0 and pH 5.40. S6
Fig. S6. Determination of the critical micelle concentration (CMC) of DMPG. S7
Fig. S7. Formation of the Dap-2PG-Ca2+ complex S8
Table S1. Assignment of the COSY signals of Dap in the fingerprint region. S9

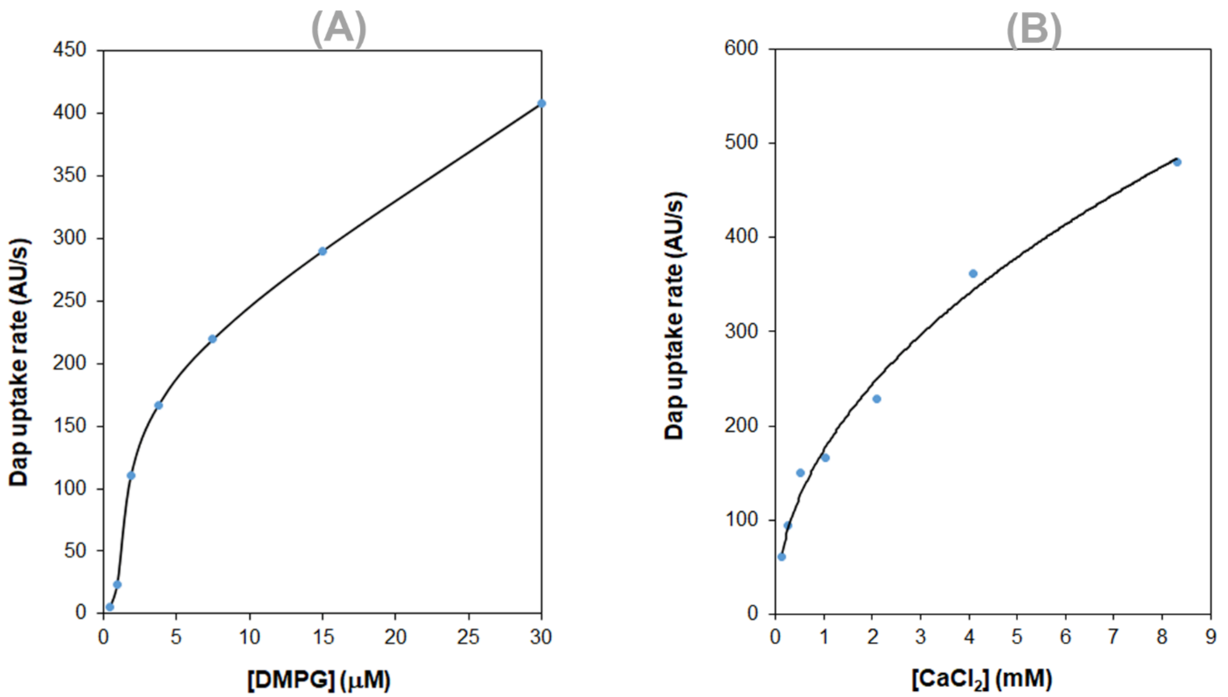


**Fig. S1.** Dependence of the Dap uptake on both DMPG and calcium. (**A**) Plot of the initial uptake rate *vs*. the DMPG content. Micelles contained 83.3 μM DMPC and DMPG at an increasing concentration; calcium concentration was 1.67 mM. (**B**) Plot of the initial Dap uptake rate *vs*. the calcium concentration. Micelles contained DMPC at 83.3 μM and DMPG at 10.8 μM. In both (A) and (B), Dap was added at 15 μM right before kinetic measurement.

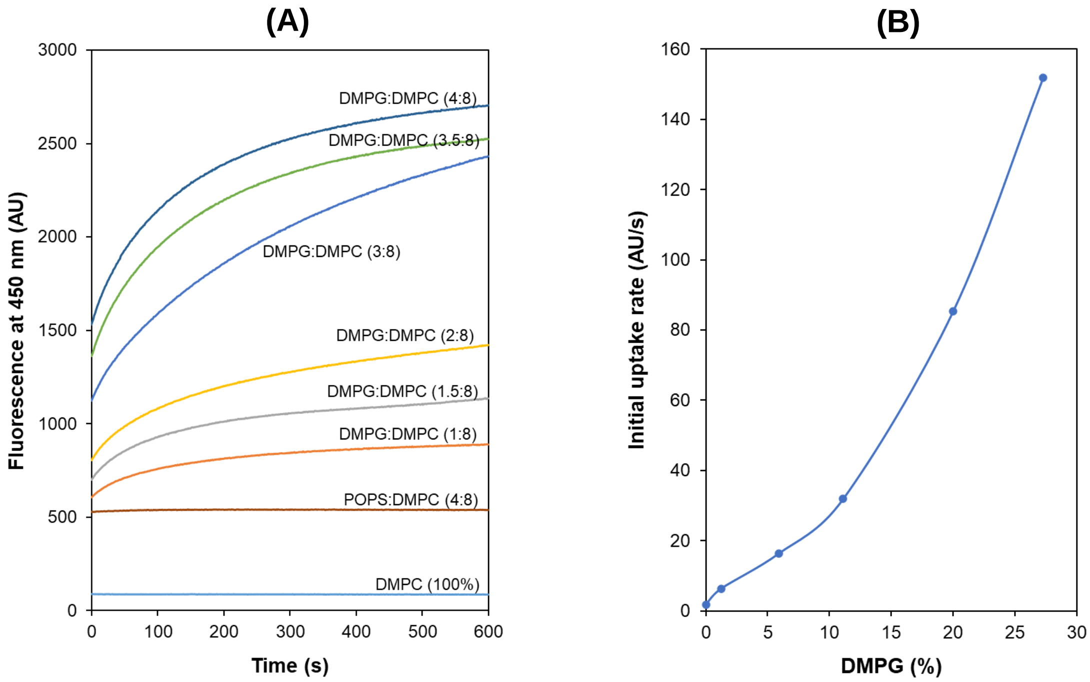


**Fig. S2**. Enhancing effect of phosphatidyl glycerol (PG) on the Dap uptake to vesicles. (A) Dose-dependent enhancement of the Dap uptake by DMPG. (B) Increase of the initial rate of Dap uptake with the membrane content of DMPG. See Experimental section for the preparation of vesicles. The uptake was measured kinetically in 20 mM HEPES (pH 7.57) using 15 µM Dap and 1.67 mM $CaCl_2$.

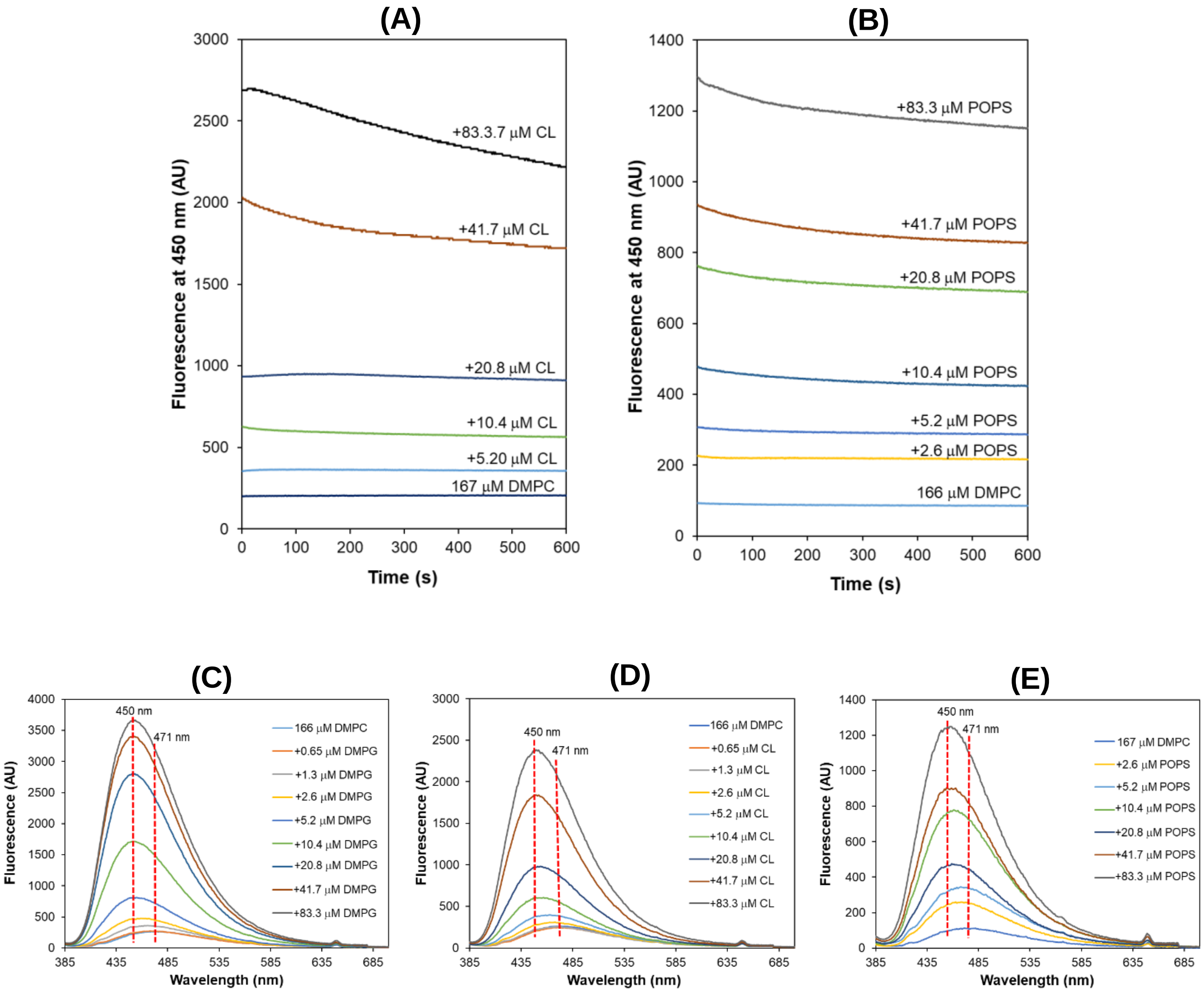


**Fig. S3.** Fluorescence enhancement and blue-shift caused by negative cardiolipin (CL) and phosphatidyserine (PS). (**A**) Increased background fluorescence but no uptake of Dap in micelles containing increasing content of CL. (**B**) Increased background fluorescence but no uptake of Dap in micelles containing increasing content of POPS. (**C**) Blue-shift of the Dap fluorescence after uptake into the DMPG-containing micelles. (**D**) Blue-shift of the Dap fluorescence after mixing with CL-containing micelles. (**E**) Blue-shift of Dap fluorescence after mixing with CL-containing micelles. All the experiments were initiated by adding Dap (final concentration = 15 µM) to 20 mM HEPES (pH 7.57) containing 166 µM DMPC, 1.67 mM $CaCl_2$ and the negative phospholipids at a varied concentration. The emission spectra in (**C**, **D** and **E**) were recorded after completing the kinetic monitoring at 450 nm.

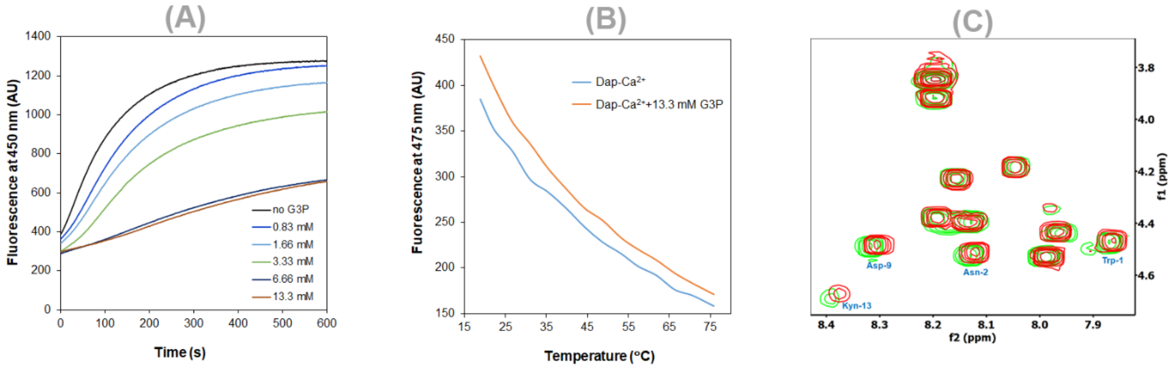


**Fig. S4**. Interaction of Dap-$Ca^{2+}$ with glycerol-3-phosphate (G3P). (**A**) G3P inhibition of the Dap uptake to DMPG-containing micelles. Dap was 15 μM and the micelles contained 167 μM DMPC, 20.8 μM DMPG in 20 mM HEPES buffer (pH 7.57) supplemented with 1.67 $CaCl_2$ and G3P at a varied concentration. (**B**) Thermal shift of the Dap fluorescence caused by G3P. The Dap (15 μM) solution contained 1.67 $CaCl_2$ with or without G3P (13.3 mM) in 20 mM HEPES (pH 7.57). (**C**) Slight changes of the Dap cross-peaks in the fingerprint region caused by G3P. The COSY spectra were recorded for 1 mM Dap in 20 mM HEPES buffer (pH/D 5.40) containing 10% $D_2O$, 1.67 $CaCl_2$ with (red) or without (green) 20 mM G3P.

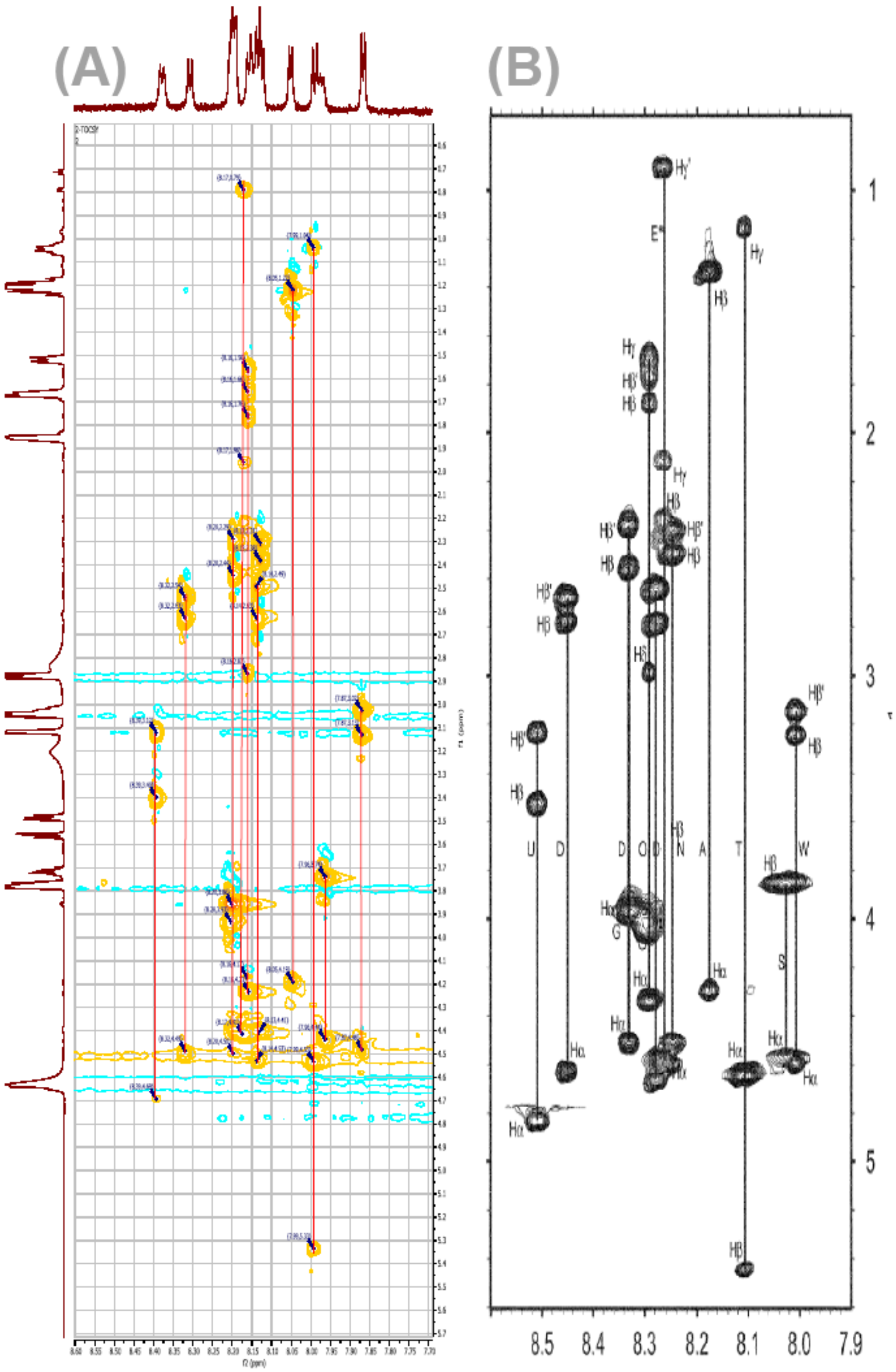


**Fig. S5**. Highly similar NαH-Hα-Hβ-Hγ spin systems in Dap at pH 5.0 and pH 5.40. (A) The NαH-Hα-Hβ-Hγ cross-peaks in the TOCSY for Dap-$Ca^{2+}$ taken in the current work at pH 5.4. (B) The corresponding NαH-Hα-Hβ-Hγ cross-peaks in TOCSY of Dap-$Ca^{2+}$ taken previously at pH 5.0 (*26*). This high level of similarity allows the assignment of the NαH-Hα cross-peaks in the fingerprint regions, as shown in Table S1, using the assignments of the previous work.

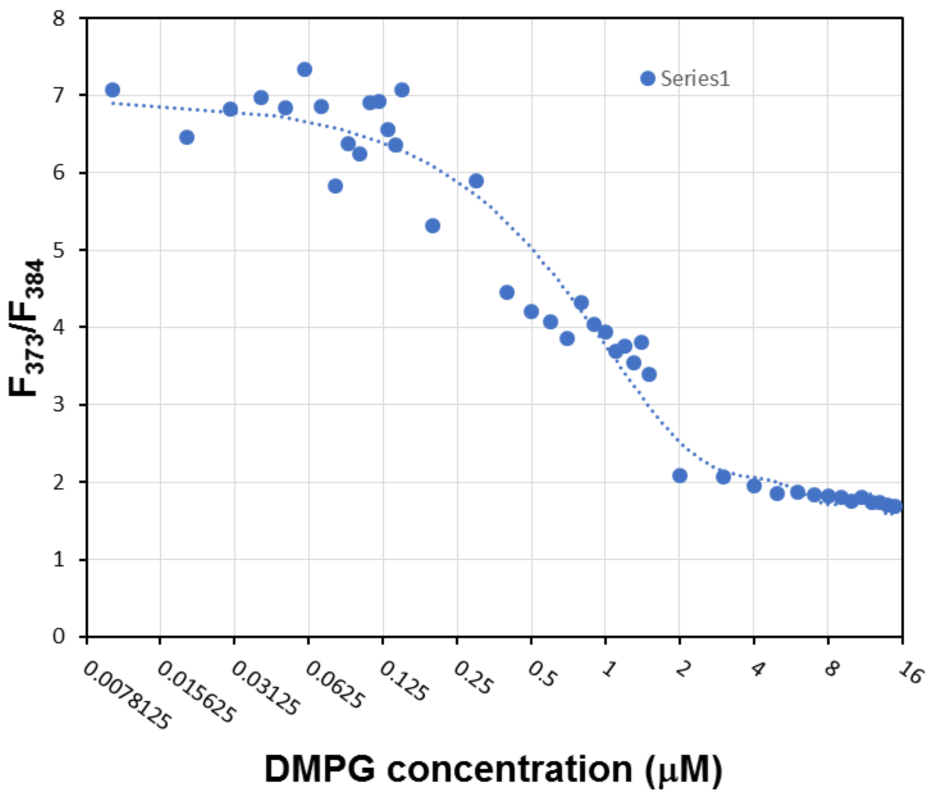


**Fig. S6**. Determination of the critical micelle concentration (CMC) of DMPG. The CMC is determined to be ~ 2.0 μM according to the relative pyrene fluorescence intensity at 373 nm and 384 nm using a reported method (*44*). The DMPG solution is thus a homogenous solution in the concentration range from 0 – 1.0 μM.

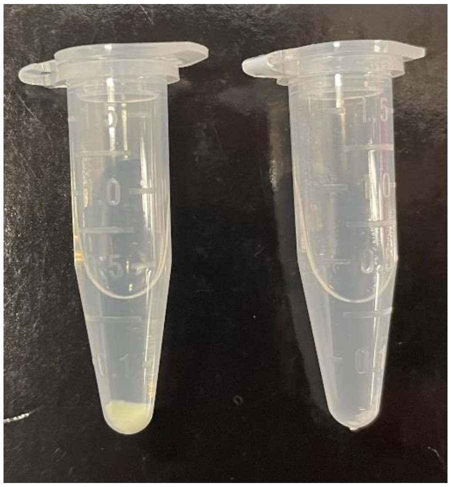

**Fig. S7**. Formation of the Dap-2PG-$Ca^{2+}$ complex. Left tube: mixing 1 mM Dap, 2 mM DMPG and 1 mM $Ca^{2+}$ to form the precipitate after centrifugation; right tube: control sample by mixing 2 mM DMPG and 1 mM $Ca^{2+}$.

**Table S1.** Assignment of the COSY signals of Dap in the fingerprint region.

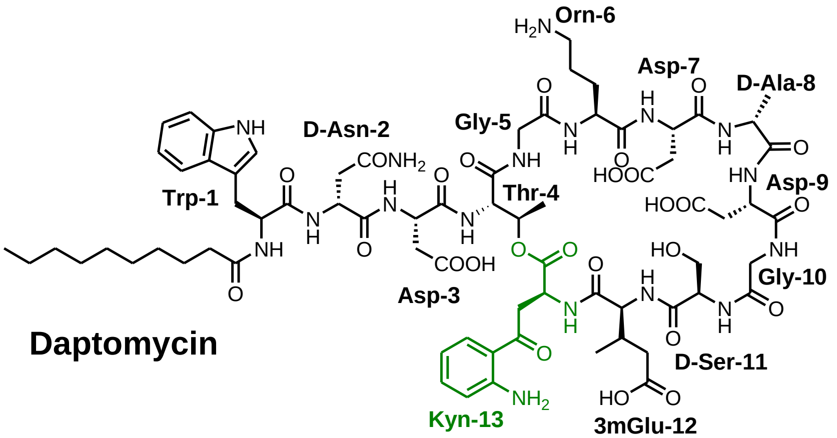


| Fingerprint COSY signals | Single Amino acid representation | Amino acid residues |
|---|---|---|
| 8.39 | U 13 | Kyn-13 |
| 8.32 | D 9 | Asp-9 |
| 8.20 | G 10, D 7 | Gly-10, Gly-5 |
| 8.16 | O 6 | Orn-6 |
| 8.14 | D 3 | Asp-3 |
| ?? Not present? | E 12 | 3mGlu-12 |
| 8.13 | N 2 | Asn 2 |
| 8.05 | A 8 | D-Ala-8 |
| 7.99 | T 4 | Thr-4 |
| 7.96 | S 11 | D-Ser-11 |
| 7.98 | ? | ? |
| 7.87 | W 1 | Trp-1 |
